# Enhanced cortical tracking of unfamiliar languages in both monolinguals and bilinguals

**DOI:** 10.64898/2026.09.21.753321

**Authors:** Qianxun Zheng, Farhin Ahmed, Aaditya Chopra, Bonnie K. Lau

**Author notes:** Corresponding author: Farhin Ahmed **Email:**, University of Washington, Box 357923, Seattle, WA 98195 USA. These authors contributed equally (Qianxun Zheng, Farhin Ahmed). **Author Contributions:** B.L. and Q.Z. designed research; Q.Z. performed research; F.A., Q.Z., A.C. and B.L. analyzed data; F.A., Q.Z. and B.L. wrote the paper. **Competing Interest Statement:** The authors declare no competing interest.

## Abstract

Humans routinely encounter speech in languages they have never heard, yet how the brain responds to such input and whether bilingual experience shapes this response remains unknown. Here, we used electroencephalography (EEG) and temporal response function (TRF) modeling to examine cortical tracking of the speech envelope in 24 English-monolingual and 24 English-Mandarin bilingual adults. Participants listened to naturally produced continuous speech in three languages: English (familiar to all), Mandarin (familiar to bilinguals only), and Vietnamese (unfamiliar to all). We report two main findings. First, both monolinguals and bilinguals showed enhanced cortical tracking for unfamiliar relative to familiar languages, evidenced by higher EEG prediction accuracy (PA). Monolinguals showed enhanced tracking for both Mandarin and Vietnamese, whereas bilinguals showed enhancement only for Vietnamese, consistent with Mandarin being a familiar language for this group. This finding suggests that enhanced cortical encoding of unfamiliar speech is a general property of the listening brain, not a signature of listening to a non-native language or reduced language proficiency. Second, bilinguals strikingly showed stronger cortical tracking than monolinguals overall, in both PA and TRF peak weights, with the TRF peak weight advantage present across all three languages, suggesting a difference in how bilingual experience shapes the neural encoding of speech. These findings have implications for understanding how the brain navigates the linguistic diversity of everyday life in an increasingly global, multilingual world.

**SIGNIFICANT STATEMENT:** More than half the world speaks more than one language, and most people regularly hear speech they cannot understand. How the brain responds to such speech, and whether bilingualism matters, is unclear. We recorded brain activity from monolingual and bilingual adults listening to familiar and unfamiliar speech. Both groups showed enhanced tracking of unfamiliar speech compared to familiar speech and bilinguals showed stronger tracking than monolinguals overall, suggesting a potential difference in how bilingual brains process speech. These findings advance our understanding of how the brain navigates speech in a multilingual world.

## INTRODUCTION

Bilingualism — the regular use of two or more languages — is the global norm (1), with more than half the world’s population estimated to speak more than one language (2). A long-standing debate concerns whether managing two languages confers cognitive and neural advantages. Some past studies have reported benefits across the lifespan: bilingual infants have shown more flexible speech learning (3), bilingual children have demonstrated enhanced metalinguistic awareness (4), and bilingual adults have shown advantages in executive functions such as inhibitory control and working memory (5, 6), along with protective effects against cognitive decline (7). However, many studies have failed to replicate these findings, reporting no differences between bilinguals and monolinguals (8–11), with inconsistencies attributed to variability in bilingual profiles, task demands, and individual differences (12–14). Crucially, much of this debate has relied on explicit behavioral measures. Neurophysiological approaches offer a complementary path by capturing brain responses directly, without requiring overt behavioral performance, and are especially suited to asking whether bilingual experience shapes the neural encoding of speech.

The human brain continuously parses incoming continuous speech to extract phonemes, words, and meaning. Cortical tracking of the speech envelope, the slow fluctuations in speech amplitude, reflects how faithfully the brain encodes ongoing speech. This tracking is measurable using electroencephalography (EEG) and quantifiable via the temporal response function (TRF) framework, which characterizes how neural activity relates to the ongoing speech signal over time. Both prediction accuracy (PA) (how well the TRF model predicts the EEG in response to the speech signal) and TRF amplitude (the magnitude of the model’s regression weights) serve as indices of cortical speech encoding fidelity. Cortical tracking is known to be modulated by attention (15–18), listening effort, and acoustic clarity (19)—but whether it is systematically shaped by language familiarity or bilingual experience, particularly for entirely unfamiliar languages, remains an open question.

A central question in bilingualism research is whether managing two languages reshapes the brain’s response to speech at a basic acoustic level, and whether that reshaping extends even to languages listeners have never encountered. TRF studies have established that cortical envelope tracking is sensitive to linguistic experience: L2 tracking is often weaker than L1 and increases as proficiency grows, suggesting that language experience influences how the brain tracks speech (20–25). However, a parallel literature suggests that reduced linguistic accessibility can also enhance envelope tracking. Non-native listeners track English speech more strongly than native listeners (24), bilingual speakers show stronger tracking for their less dominant than dominant language (25), and phonotactically familiar but uncomprehended speech can elicit greater acoustic tracking than understood speech (26). Together, these findings suggest that enhanced envelope tracking emerges not only from language experience, but also when speech is less linguistically accessible. This is consistent with the view that the brain falls back on low-level acoustic cues when higher-level linguistic representations are unavailable.

This raises a question that neither line of work can answer alone. Is this enhanced acoustic tracking a property of the listener — does the bilingual brain, shaped by lifelong experience of navigating competing linguistic systems, show a distinct neural response even to languages it has never encountered? Or is it a property of the situation — a universal consequence of linguistic inaccessibility that any listener would show, regardless of their language background? The two lines of work have never been brought together in a design that can distinguish these accounts: studies comparing monolinguals and bilinguals have always done so using languages already known to at least one group, while studies of unfamiliar or incomprehensible speech have not included a bilingual comparison. Answering the question requires testing monolinguals and bilinguals on a language equally unfamiliar to both. Only one study included a language that is equally unfamiliar to both monolinguals and bilinguals. Van den Eynde et al. (27) tested Dutch monolingual and Dutch-French bilingual children on Dutch, French, and Italian, and found greater tracking on comprehensible language than incomprehensible language for both groups. This finding leaves open whether similar effects extend to adults and to other combinations of familiar and unfamiliar languages.

To address this gap, we recorded EEG from 24 English-monolingual and 24 English–Mandarin bilingual adults while they listened to naturally produced continuous speech in three languages: English, Mandarin, and Vietnamese. English was familiar to both groups; Mandarin was familiar to bilinguals only; and Vietnamese was entirely unfamiliar to all participants, with none reporting any prior exposure. Using TRF modeling, we quantified both PA and TRF amplitude as indices of cortical speech tracking across listener groups and languages. This design allows us to disentangle the effect of language familiarity from that of bilingual experience and to ask, whether the bilingual brain encodes an entirely novel language differently from the monolingual brain.

## RESULTS

EEG data were recorded from 24 English monolinguals and 24 English–Mandarin bilinguals as they listened to continuous speech in 3 language conditions: English, Mandarin, and Vietnamese (Fig. 1A). All participants reported no prior exposure to Vietnamese. To quantify cortical speech tracking, we used a forward encoding model that relates the speech envelope to the recorded EEG, implemented with the multivariate Temporal Response Function (mTRF) toolbox (28, 29). TRF models were trained and tested using a leave-one-out cross-validation approach. The resulting models were used to predict the EEG signal in response to speech across different languages (Fig. 1B; see Methods for details). Model performance was quantified as Pearson’s correlation between the actual and the predicted EEG, hereafter referred to as PA.

**Figure 1.**
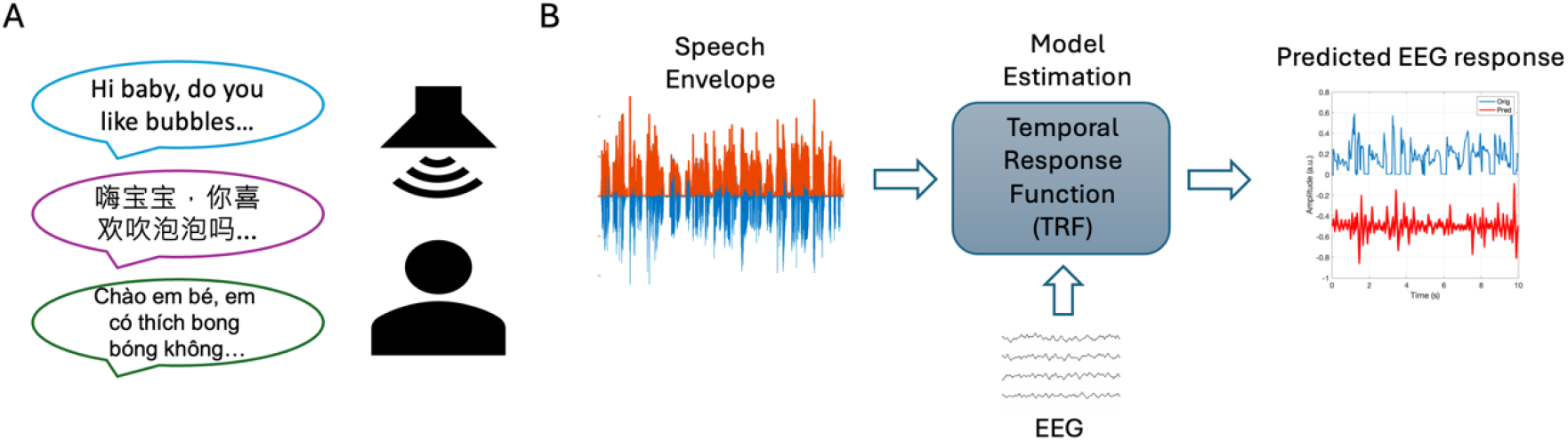
(A) Experimental set up. Stimuli consist of naturally produced speech in English, Mandarin and Vietnamese. Participants listened to ten 1-minute trials for each language condition presented from a speaker at 0° azimuth at 70 dB SPL with order and condition randomized. (B) Schematic diagram of the forward/encoding TRF model. The encoding model predicts the EEG signal from the envelope of the stimuli.

Figure 2 shows the PA, averaged across participants and all EEG channels. To test the effect of listener group and language condition on PA, we fit an LME model including group (monolingual, bilingual), condition (English, Mandarin, Vietnamese) and their interaction as fixed factors and random intercepts for participants. The model revealed a significant main effect of group (*F*(1,46) = 12.03, *p = 0*.*001, η*^*2*^*p = 0*.*207)*, condition (*F*(2,92) = 34.91, *p < 0*.*001, η*^*2*^*p = 0*.*431)*, and a significant interaction between group and condition (*F*(2,92) = 3.48, *p = 0*.*035, η*^*2*^*p = 0*.*070)*, suggesting that the effect of language on cortical speech tracking differs between monolingual and bilingual listeners. Thus, post-hoc comparisons were conducted to further evaluate the differences, with False Discovery Rate (FDR) corrections applied for multiple comparisons.

**Figure 2.**
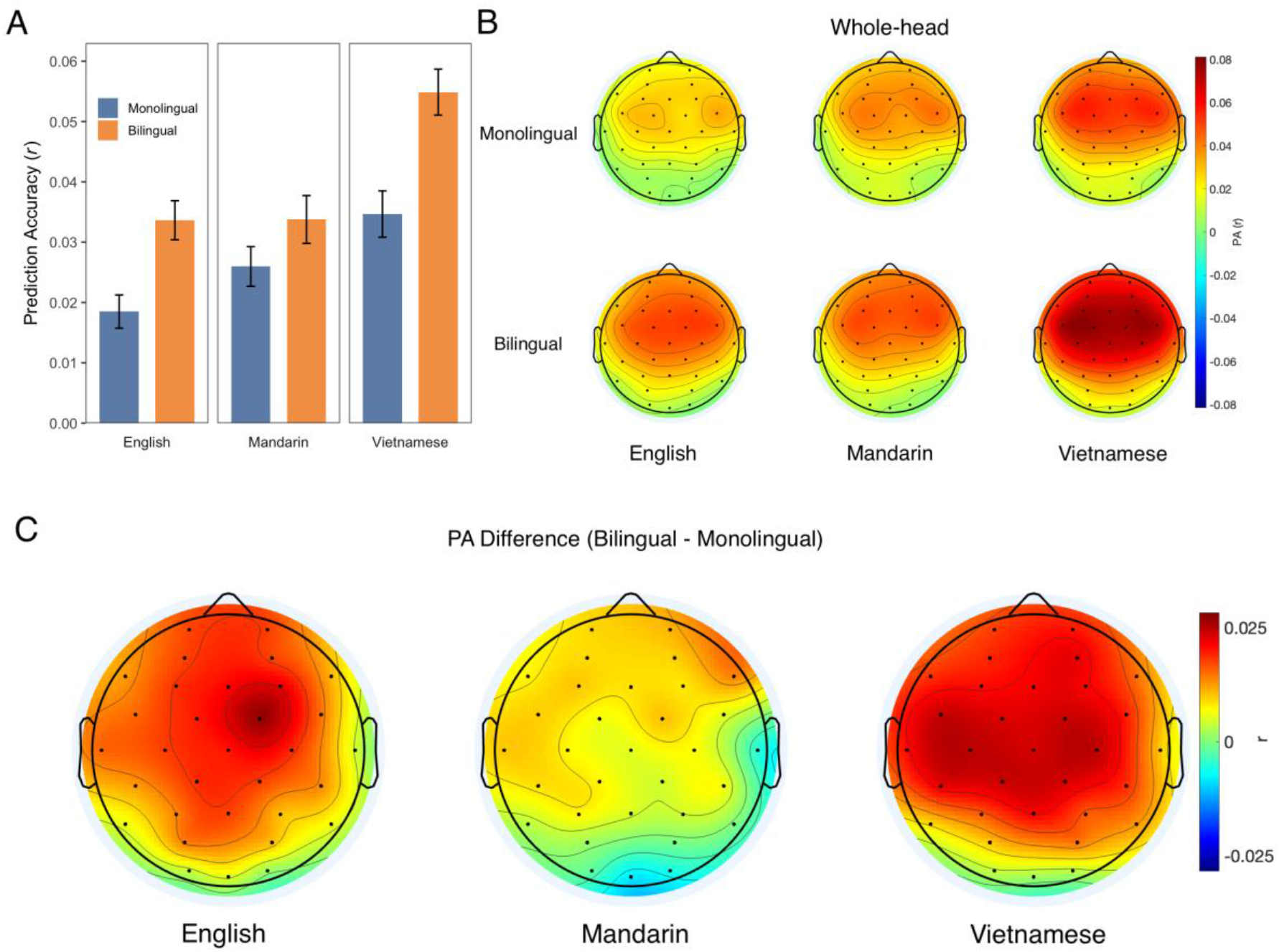
(A) Whole head PA (r) in all three conditions for monolinguals and bilinguals averaged across all EEG channels (mean ± SEM). (B) Topographical distribution of PA across conditions in monolinguals (top) and bilinguals (bottom). Red indicates higher PA and blue indicates lower PA. (C) Topographic distribution of PA difference between monolinguals and bilinguals for each condition, where red indicates greater difference and blue indicates less difference.

### Both monolinguals and bilinguals show stronger cortical tracking in unfamiliar language

We first examined how PA varied within each listener group across language conditions. For both monolinguals and bilinguals, PA was significantly higher in unfamiliar than familiar languages. Monolinguals showed significantly higher PA for both Mandarin and Vietnamese than for English (English vs Mandarin: *t =* -2.235, *p* = 0.028; English vs Vietnamese: *t* = -4.832, *p* < 0.001). Similarly, bilinguals showed significantly higher PA for Vietnamese, the unfamiliar language, than for either English or Mandarin (Vietnamese vs English, *t =* 6.348, *p* < 0.001; Vietnamese vs Mandarin, *t =* 6.306, *p* < 0.001), whereas PA did not differ between English and Mandarin, their familiar languages (*t =* -0.042, *p* = 0.967). Scalp topographies showed a fronto-central distribution of PA across groups and conditions (Fig. 2B), consistent with the canonical scalp distribution of auditory EEG responses to continuous speech (Fig. 2B). This fronto-central pattern was most pronounced for Vietnamese, particularly in bilinguals, mirroring the condition effects observed in the channel-averaged PA results. Together, these findings suggest that cortical tracking was strongest for unfamiliar languages – Mandarin and Vietnamese in monolinguals, and Vietnamese in bilinguals.

### Bilinguals show stronger cortical tracking than monolinguals in English and Vietnamese

We next examined how PA differed between listener groups within each language condition using separate linear regression models with PA as the outcome and listener group as the predictor.

Bilinguals showed significantly higher PA than monolinguals for English (*t* = 3.56, *p* < 0.001) and Vietnamese (*t =* 3.73, *p* < 0.001), but not Mandarin (*t* = 1.52, *p* = 0.137). The topographic distribution of group differences (bilingual PA − monolingual PA) further showed clear group differences for Vietnamese and English, with weaker difference for Mandarin (Fig. 1C), consistent with the channel-averaged PA results. Together, these findings indicate that bilinguals showed stronger cortical tracking than monolinguals for English and Vietnamese — languages that were familiar and unfamiliar, respectively, to both groups.

### Bilinguals show larger TRF peak weights than monolinguals

We next examined the TRF model weights to characterize the temporal and spatial dynamics of cortical speech tracking. TRF weights reflect the estimated EEG response to speech envelope at different time lags. For example, TRF weights at a lag of 100 ms reflect the relationship between the speech envelope at time *t* and the EEG response at time *t* +100 ms. We summarized these dynamics by averaging the TRF weights within frontal, temporal, occipital, and left- and right-hemispheric region-of-interests (ROIs). Figure 3A and B show the TRF time courses across ROIs, and Figure 3C and D show their scalp distributions from 0 to 300 ms.

**Figure 3.**
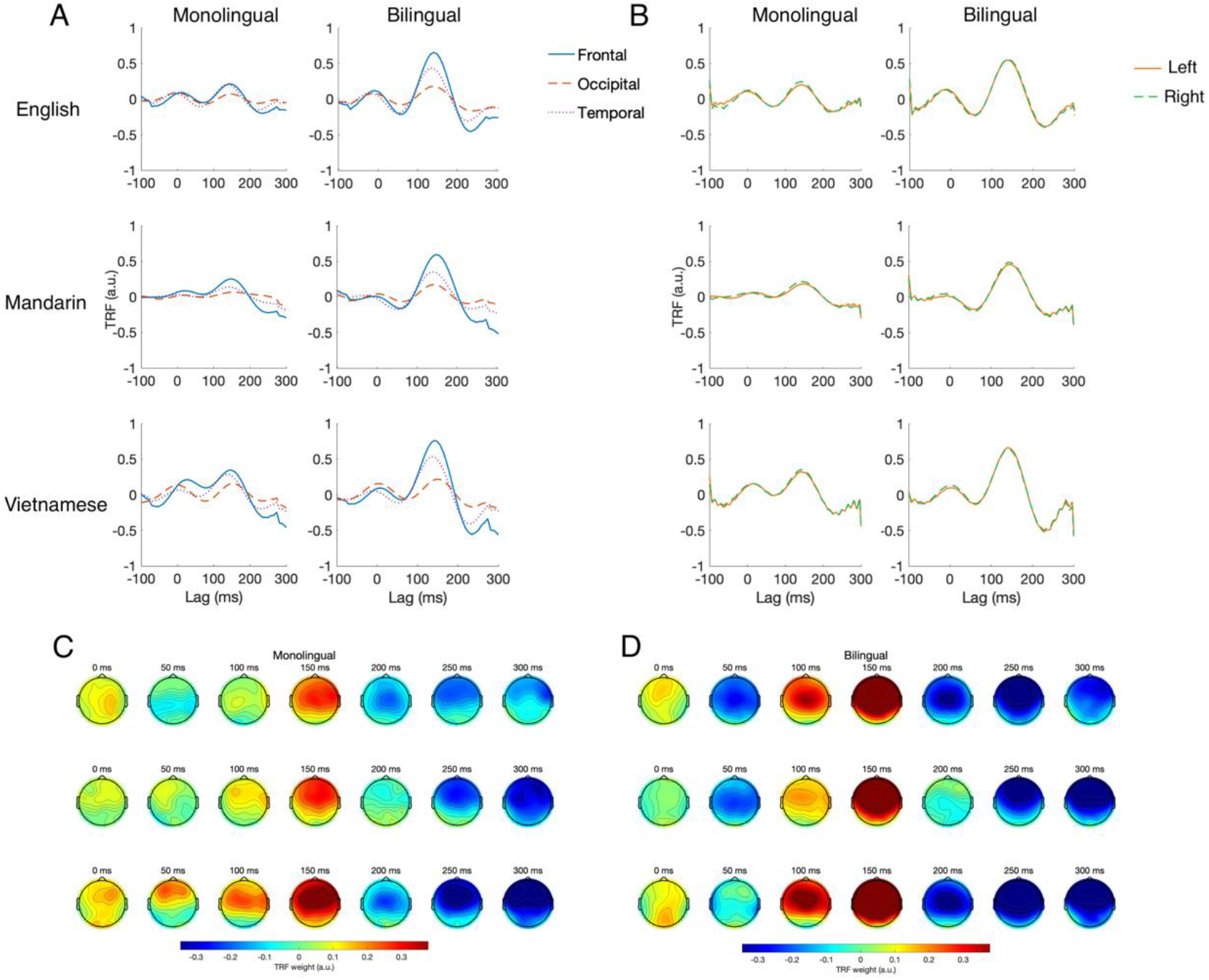
TRF weights in monolinguals and bilinguals averaged across subjects at (A) frontal, occipital, and temporal ROIs and at (B) left- and right-hemispheric ROIs in all three language conditions. Topographic distribution of TRF weights across different time lags for monolinguals (C) and bilinguals (D), where red indicates higher weights and blue indicates less weights.

**Figure 4.**
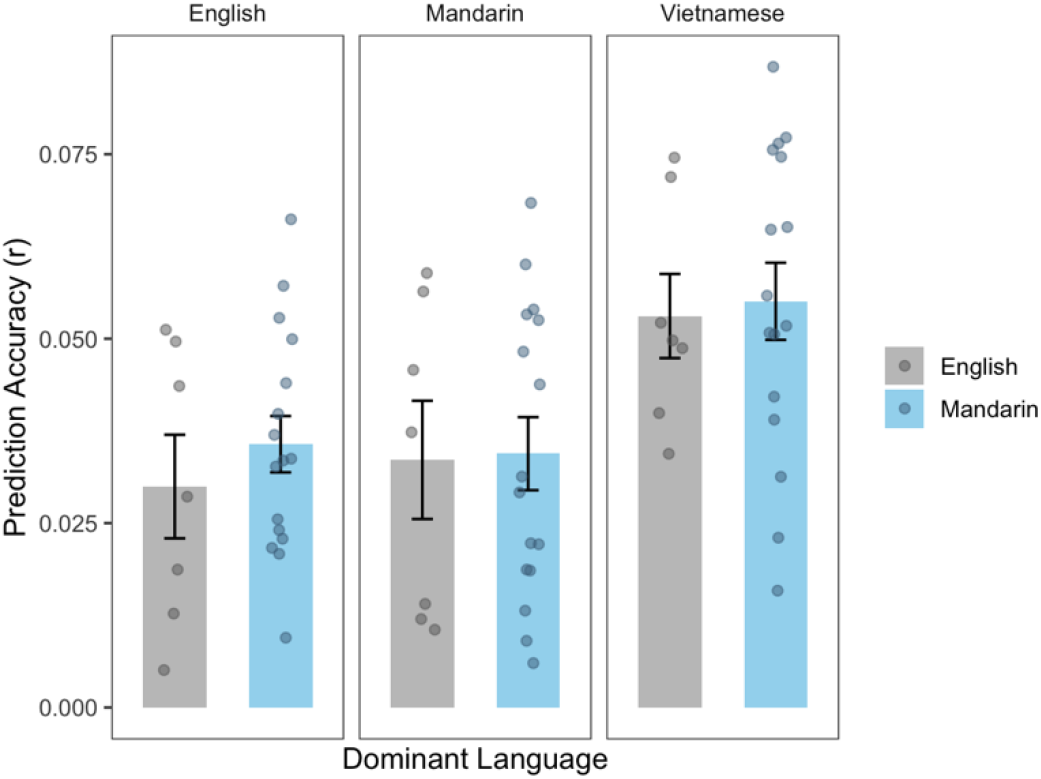
PA of bilinguals across 3 conditions (mean ± SEM) as a function of language dominance, defined as the language with the higher self-rated proficiency score on LEAP-Q. Gray indicates English as the dominant language, whereas blue indicates Mandarin as the dominant language.

Across groups and language conditions, TRF weights showed a similar overall morphology, with a prominent positive peak around 150 ms. TRF weights were largest over frontal channels, followed by temporal channels, and smallest over occipital channels (Fig. 3A), consistent with canonical TRF responses to continuous speech (30). Left- and right-hemisphere ROIs showed highly similar waveforms and amplitudes, suggesting no clear hemispheric asymmetry in the TRF weights (Fig. 3B). The scalp topographies showed a similar pattern, with the strongest TRF weights emerging around 150 ms and a fronto-central distribution across groups and language conditions (Fig. 3C, D). Importantly, bilinguals showed larger TRF weights around this peak than monolinguals. To statistically evaluate this group difference, peak weights within the 100–200 ms post-stimulus window were analyzed using an LME model including Group, Condition, ROI, and their interactions as fixed effects and random intercepts for subjects. The model revealed a significant main effect of Group (*F*(1,59.07) = 18.53, *p* < 0.0001, *η*^*2*^*p* = 0.24) with bilinguals exhibiting larger TRF peak weights than monolinguals. Follow-up estimated marginal mean contrasts confirmed this pattern across all language conditions (Bilingual vs Monolingual: English, *t*(66.9) = 4.15, *p* = 0.0001; Mandarin, *t*(66.9) = 3.31, *p* = 0.0015; and Vietnamese, *t*(71.1) = 4.82, *p* < 0.0001). These findings indicate that, overall, the speech envelope elicited stronger cortical responses in bilinguals across languages.

### Bilingual language experience did not significantly modulate cortical speech tracking within the bilingual group

Given the inherently heterogenous nature of bilingual language background, we examined whether variability within the bilingual group modulates cortical tracking of speech. Bilingual participants completed the Language Experience and Proficiency Questionnaire (LEAP-Q) which measures self-reported language exposure and proficiency (31). We characterized each bilingual participant along three dimensions of language background: current language exposure, spoken language proficiency, and language dominance. Participants varied in current language exposure (English: M = 62.79%, SD = 18.03%; Mandarin: M = 38.13%, SD = 18.87%) and in self-rated spoken language understanding on a 10-point scale, with higher scores indicating greater understanding (English: M = 7.92, SD = 2.06; Mandarin: M = 9.13, SD = 1.23). We categorized bilinguals as English- or Mandarin-dominant based on which language they reported higher proficiency in.

Language dominance did not significantly predict cortical tracking: a linear mixed-effects model with PA as the outcome and language dominance, condition, and their interaction as predictors revealed no significant main effect of dominance (*F*(1,22) = 0.13, *p* = 0.718, *η*^*2*^*p* = 0.006). We further examined whether current language exposure and proficiency were associated with PA using Pearson correlations. Neither English exposure, Mandarin exposure, English understanding score, nor Mandarin understanding score was significantly correlated with PA in any language condition (all *p* > 0.05). These findings suggest that the bilingual advantage in cortical tracking observed here is not driven by individual differences in language dominance, exposure, or proficiency within the bilingual group.

## DISCUSSION

This study examined how monolingual and bilingual adults respond to speech across familiar and unfamiliar languages. Two main findings emerged. First, both groups showed enhanced cortical tracking for their respective unfamiliar language, demonstrating for the first time that enhanced acoustic-level encoding of unfamiliar speech is a general property of the human listening brain — not a signature of non-nativeness or reduced proficiency, as previously proposed. This resolves a fundamental ambiguity in existing literature: prior studies comparing monolinguals and bilinguals used only languages familiar to at least one group, while studies using unfamiliar speech lacked a bilingual comparison, leaving it unclear whether enhanced tracking reflected the listener’s background or the listening situation itself. Second, bilinguals showed greater cortical tracking than monolinguals overall, in both PA and TRF peak weights, regardless of language familiarity. This suggests that bilingual experience confers a familiarity-independent enhancement in the strength of the neural response to speech. Within the bilingual group, cortical tracking did not vary significantly with language dominance or self-reported language exposure.

In our study, English monolinguals showed stronger cortical tracking for Mandarin and Vietnamese than English, whereas English-Mandarin bilinguals showed stronger tracking for Vietnamese than both English and Mandarin. At first glance, enhanced cortical tracking for unfamiliar languages may seem surprising, given that listeners had no access to the linguistic content. Therefore, a central consideration in interpreting these findings is what cortical speech tracking indexes. Our analysis is based on how well EEG tracks the amplitude envelope of the acoustic speech. This measure has been widely used in speech-neuroscience research (32–36). The speech envelope captures slow fluctuations in acoustic energy that carry temporal information important for parsing continuous speech, that broadly align with syllabic and phrasal timescales (37, 38). These low-frequency acoustic cues are thought to support segmentation of continuous speech into units to be used by later stages of speech processing. As such, envelope tracking is not a direct measure of speech comprehension. Rather, it reflects acoustic-temporal encoding that can be further modulated by higher-level cognitive and linguistic factors, including attention (15, 39) and speech intelligibility (40, 41). Thus, enhanced tracking for unfamiliar speech may reflect a shift toward reliance on acoustic cues, rather than paradoxical processing efficiency.

Two non-exclusive interpretations may explain this unfamiliar-language effect. First, unfamiliar speech may have drawn greater attention: attention modulates cortical speech tracking, and listeners attending one speech stream show stronger cortical responses to it than to an ignored stream (15, 16, 18, 34, 39, 42). Sustained attention is similarly associated with stronger tracking (43), though because attention was not directly manipulated in the present study, this interpretation remains tentative. Important to consider nonetheless is that attention need not be directed. Some speech sounds may be more acoustically salient than others and may therefore draw a listener’s attention in a bottom-up stimulus-driven manner. For example, infant-directed speech, which is characterized by exaggerated prosody and slower tempo, elicits stronger cortical tracking than adult-directed speech in infants (44), even though infants have immature attentional mechanisms. In our study, unfamiliar languages being novel, may have been acoustically salient to listeners, increasing attention and thereby enhancing cortical tracking.

Second, stronger envelope tracking for unfamiliar languages may reflect greater reliance on low-level acoustic cues when higher-level linguistic representations are unavailable, consistent with evidence that acoustic-level encoding is enhanced for less familiar languages while phonetic, lexical, and semantic features are more strongly encoded for familiar ones (25, 26, 45, 46). What neither account resolves, however, is whether this enhanced tracking is specific to listener types or arises in any listener confronted with an unfamiliar language.

Our design allows us to directly resolve this ambiguity, and in doing so, reframe the contributions of three prior studies. Reetzke et al. (24) reported enhanced envelope tracking in non-native (Mandarin-English bilingual) listeners relative to native English listeners and interpreted this as reflecting increased reliance on acoustic cues in non-native listening. However, their design could not determine whether this enhancement was specific to non-native listeners or would arise in any listener confronted with a linguistically inaccessible language. Our data resolve this: English monolinguals showed enhanced tracking for both Mandarin and Vietnamese — languages unfamiliar to them — demonstrating that heightened envelope tracking of unfamiliar speech is a property of the listening situation, not a signature of being a non-native listener. Tezcan et al. (26) reached a related conclusion using Dutch listeners hearing acoustically familiar but incomprehensible French speech. Our Vietnamese condition extends this to a language that is phonotactically and prosodically novel, ruling out acoustic familiarity as an explanation and establishing enhanced unfamiliar-language tracking as a truly general property of the human auditory brain. Finally, Pérez-Navarro et al. (25) showed that in bilingual children, the less dominant language elicits stronger acoustic-level tracking than the dominant language. Our TRF peak weight results extend this finding to adults: bilinguals showed significantly larger TRF peak weights than monolinguals across all three language conditions, including Mandarin, which was a familiar language for bilinguals. This suggests that the bilingual advantage in how strongly the brain weighs the speech envelope is not contingent on language familiarity but may reflect a more fundamental difference in how the bilingual auditory system encodes incoming speech.

Our findings can be compared with a prior study that used a similar multilingual design in monolingual and bilingual children (27) but reported a different pattern. According to their study, cortical tracking was stronger for familiar language compared to unfamiliar language in both monolingual and bilinguals. At first glance, this finding appears to contrast with our observation. However, several methodological differences may account for this discrepancy. First, although both studies tested a third unfamiliar language, their set of languages (Dutch, French, and Italian) are all Indo-European languages that share many phonological and rhythmic properties. Whereas we included languages that are more acoustically distinct from participants’ known language.

Second, their study tested children aged 6-8 years, whereas our participants were adults, raising the possibility that the effect of language familiarity on cortical tracking differs across development. Finally, their task required active listening through story-related images and comprehension questions, whereas our study employed a passive listening paradigm. As attention is known to modulate cortical tracking, top-down attention may enhance cortical tracking of comprehensible speech. In contrast, passive listening draws attention from bottom-up.

Together, these differences suggest that the relationship between language familiarity and cortical tracking may depend on the linguistic, developmental, and listening context. The absence of a group difference in PA for Mandarin is noteworthy. Mandarin placed the two groups in opposite familiarity positions — familiar to bilinguals, unfamiliar to monolinguals, such that opposing familiarity effects are likely cancelled at the group level. Notably, bilinguals showed larger TRF peak weights than monolinguals even for Mandarin (*p* = 0.0015), a pattern that dissociates from the PA results and carries distinct mechanistic significance. PA reflects how precisely the brain tracks the temporal speech envelope, whereas TRF peak weights reflect how strongly a unit change in the speech envelope drives EEG activity at the peak response latency. For Mandarin, a familiarity-driven boost in monolinguals’ PA narrows the group difference in tracking fidelity, but TRF peak weights, being less sensitive to this ceiling effect, still reveal a robust bilingual advantage. This advantage peaked around 150 ms with a fronto-central scalp distribution, consistent with generators in auditory cortex and adjacent prefrontal regions implicated in top-down modulation of speech processing. Together, these findings suggest that bilingual experience strengthens the neural response to the speech envelope itself, a familiarity-independent enhancement that PA alone does not reveal.

One question that arises in bilingual research concerns how bilingualism is defined. We employed a broad definition, requiring a self-reported comprehension score above 3 on a 10-point scale from the LEAP-Q for both languages, where 10 being perfect understanding of spoken language (31). This criterion was selected as most of our bilingual participants are native Mandarin speakers enrolled in college, indicating that they had already reached at least working proficiency in English prior to college admission. Indeed, cortical tracking did not vary significantly with current language exposure, spoken language proficiency, or language dominance language dominance within the bilingual group. This suggests that, within the bilingual group, the unfamiliar-language effect was not strongly explained by individual differences in language dominance, exposure, or comprehension of the familiar languages. As such, our findings should be interpreted as reflecting attention-related modulation and/or greater reliance on acoustic-level speech encoding, rather than differences in comprehension of the languages tested.

Taken together, our study offers insight into how language experience and language familiarity shape the brain’s responses to speech. Several future directions follow from these findings. Because Mandarin and Vietnamese are both tonal and syllable-timed languages, whereas English is non-tonal and stress-timed, future work should examine whether pitch and rhythmic structure contribute to the enhanced tracking observed for Vietnamese, and whether bilinguals’ prior experience with Mandarin supports sensitivity to acoustic features shared with Vietnamese. The TRF framework can also be extended to higher-level linguistic features — phonetic, lexical, and semantic representations — to determine whether the bilingual enhancement observed here is specific to acoustic-level encoding or extends further up the processing hierarchy (47–51). Finally, examining these effects in bilingual infants and children could clarify whether bilingual advantages in neural speech encoding emerge early and whether they shape novel language learning across development.

## MATERIALS AND METHODS

### Participants

Participants were 24 English monolinguals (*M* = 21.7 years, *SD* = 4.18, 18 female) and 24 English-Mandarin bilinguals (*M* = 21.5 years, *SD* = 2.32, 19 female) with high proficiency in both English (Self-rated understanding score: *M* = 7.92, *SD* = 2.06), and Mandarin (Self-rated understanding score: *M* = 9.13, *SD* =1.23). Participants were recruited from research subject pool and flyers posted around campus. All participants reported no previous exposure to Vietnamese, no history of hearing loss, and no other health or neurodevelopmental concerns that would interfere with testing. Participant race was self-reported. English monolinguals included 16 White, 3 Asian, 2 Black/African American, and 1 native Hawaiian/Pacific Islander, 1 More Than One Race, and 1 other race. All 24 English-Mandarin bilinguals identified as Asian. All participants completed the Language Experience and Proficiency Questionnaire (LEAP-Q), a self-report tool designed to assess participants’ language background, usage, and self-rated proficiency across multiple domains, such as speaking and understanding (31). Bilingualism was defined as scoring higher than 3 out of 10 in language comprehension for both English and Mandarin on the LEAPQ. On the day of testing, all participants were required to pass a tympanometric screen and an audiometric screen (< 20 dB HL) at octave frequencies between 500 Hz and 8000 Hz.

All protocols were conducted according to protocols approved by the Institutional Review Board at the University of Washington. All participants provided written consent prior to testing, and either received course credit or monetary compensation for their time.

### Stimuli

The stimuli consisted of naturally recorded infant directed speech (IDS) in three language conditions: 1) English, 2) Mandarin, and 3) Vietnamese. IDS was chosen for two reasons. First, its acoustic properties, including higher pitch, slower speaking rate, and exaggerated prosody, enhance acoustic salience and sustain listener’s attention, which supports robust cortical tracking of the speech envelope in adult listeners (44). Second, using IDS enables direct comparison with planned infant data, allowing the developmental trajectory of language-familiarity and bilingual effects to be examined within the same paradigm. The IDS stimuli were recorded by female native speakers of each language. English and Mandarin stimuli were each recorded by two different native language speakers. Vietnamese stimuli were recorded by one native language speaker. Stimulus recording took place in a sound-attenuated booth with the speakers seated 25 cm from a microphone. Each speaker recorded fifteen to thirty minutes of infant directed speech, and from which five 1-minute segments were manually selected from each English and Mandarin speaker, and ten 1-minute segments were selected from the Vietnamese speaker. All stimuli were inspected for artifacts and then RMS normalized in Praat (52). Silent gaps in the speech exceeding 300 ms were reduced to 300 ms using a custom MATLAB script. The final stimuli consist of ten 1-minute trials for each language adding up to a total recording time of 30 minutes.

### EEG Data Acquisition

Participants completed a passive listening EEG recording that took place in a sound attenuated booth. EEG data were recorded using a 32-channel BioSemi Active Two system at a sampling rate of 2048 Hz. During the recording, participants sat in a chair and watched a silent movie with subtitles. The stimuli were presented at an overall level of 70 dB SPL via a speaker placed at 0° azimuth, approximately 1 meter in front of the subject with randomized condition and trial order. All participants completed the full experiment.

### EEG Data Analysis

For individual datasets to be included in the analysis, the dataset must have less than 20% noisy channels (< 6 electrodes) and the TRF model must achieve PA greater than chance.

### Pre-preprocessing

Raw EEG data were processed offline using the EEGLAB toolbox in MATLAB (53) according to the following steps. First, EEG data were epoched into 1-minute segments, aligned with the onset of the stimulus in each trial. A second-order Butterworth band-pass filter with cutoff frequencies of 1-8 Hz were then applied. To remove artifact, we applied artifact subspace reconstruction (ASR), as in previous TRF studies (44, 54). ASR first identified the cleanest segment of the recording and used this segment to estimate baseline statistics. The full dataset was then processed using a 500-ms sliding-window subspace decomposition to identify components with abnormally high variance relative to the clean baseline data, using a threshold of 20 SDs. Channels contributing to these high-variance subspaces were reconstructed using a mixing matrix derived from the clean baseline data. Following ASR, channels with excessive residual noise were identified based on channel-wise kurtosis, with rejection thresholds set at 3 SDs from the mean using the EEGLAB function pop_rejchan. Spherical interpolation was then performed to interpolate the noisy channels. Finally, the data were down sampled to 128 Hz to reduce processing time and rereferenced to the average of the left and right mastoid electrodes.

### Multivariate Temporal Response Function (mTRF) Analysis

To quantify cortical tracking of speech, we used the mTRF approach (28, 29), which models the relationship between speech features and the EEG response at each channel over a range of time lags. For the speech feature, here, we focus on the envelope. The amplitude envelope of each speech stimulus was extracted using the Hilbert transform and down sampled to 128 Hz to match the EEG sampling rate. We then fit forward TRF models over a time-lag window from -100 to 300 ms. Model weights were estimated by minimizing the mean squared error between the actual and predicted EEG responses using ridge regression: TRF = (S^T^S + λI)^-1^S^T^R; where λ is the ridge regression parameter that controls overfitting, I is the identity matrix, S(t) and R(t) are the lagged time series of the stimulus envelope and the EEG response respectively. The ridge parameter λ was optimized using leave-one-out cross-validation. For each subject and condition, the model was trained on 9 out of 10 trials and tested on the held-out trial, repeating the procedure until each trial served as the test set once. The λ value yielding the highest Pearson correlation between predicted and actual EEG responses, averaged across trials and channels, was selected. Pearson correlation at the optimal λ was reported as PA.

To determine whether TRF model performance exceeded chance, we used a nonparametric permutation-based approach. For each permutation, the true stimulus–EEG correspondence was disrupted by applying a random circular shift to the stimulus. The same cross-validation procedure described above was then used to compute PA for the shuffled data. This process was repeated 100 times for each subject and condition to generate a null distribution of chance-level PA. Wilcoxon-signed rank tests determined that PAs were significantly above change for both monolinguals and bilinguals in all listening conditions (Monolinguals: *z* = 3.77, *p* < 0.001, English; *z* = 4.17, *p* < 0.001, Mandarin; *z* = 4.26, *p* < 0.001, Vietnamese; Bilinguals: *z* = 4.29, *p* < 0.001, English; *z* = 4.29, *p* < 0.001, Mandarin; *z* = 4.29, *p* < 0.001, Vietnamese).

### Statistical Analysis

To examine effects of group and condition on PA, we fit a linear mixed-effects model (LME) with group (monolinguals, bilinguals), condition (English, Mandarin, Vietnamese), and their interaction as fixed effects and a random intercept to account for across-subject variability. Fixed effects were assessed using Type III ANOVA with Satterthwaite approximation, and post-hoc pairwise comparisons were corrected using the False Discovery Rate (FDR; α = 0.05). To examine group differences within each language condition, we additionally fit separate linear regression models with PA as the outcome and listener group as the predictor.

To assess whether language background predicted PA within the bilingual group, we fit a separate LME with language dominance (English-dominant, Mandarin-dominant), condition, and their interaction as fixed effects and a random intercept. We additionally computed Pearson correlations between PA and four language background measures: English exposure, Mandarin exposure, English understanding score, and Mandarin understanding score. All analyses were conducted in R version 4.5.1 within RStudio (55) using lmerTest version 3.1.3 and lme4 version 1.1.37.

## ACKNOWLEDGEMENTS

This research was supported by T32DC005361-21 to F.A. We thank participants for their contributions to this research.

## DATA AVAILABILITY

Data are available from the corresponding author upon request

## Notes

### Competing Interest Statement

The authors have declared no competing interest.

## REFERENCES

1. L. Wei, Ed., The Bilingualism Reader, 0 Ed. (Routledge, 2003).

2. F. Grosjean, Bilingual: Life and Reality (Harvard University Press, 2010).

3. Á. M. Kovács, J. Mehler, Flexible learning of multiple speech structures in bilingual infants. Science 325, 611–612 (2009).

4. J. Cromdal, Childhood bilingualism and metalinguistic skills: Analysis and control in young Swedish–English bilinguals. Applied Psycholinguistics 20, 1–20 (1999).

5. O. A. Olulade, et al., Neuroanatomical Evidence in Support of the Bilingual Advantage Theory. Cereb Cortex 26, 3196–3204 (2016).

6. A. Miyake, et al., The unity and diversity of executive functions and their contributions to complex “frontal lobe” tasks: a latent variable analysis. Cognitive Psychology 41, 49–100 (2000).

7. E. Bialystok, F. I. M. Craik, M. Freedman, Bilingualism as a protection against the onset of symptoms of dementia. Neuropsychologia 45, 459–464 (2007).

8. E. Antón, Y. Fernández García, M. Carreiras, J. A. Duñabeitia, Does bilingualism shape inhibitory control in the elderly? Journal of Memory and Language 90, 147–160 (2016).

9. E. Antón, M. Carreiras, J. A. Duñabeitia, The impact of bilingualism on executive functions and working memory in young adults. PLOS ONE 14, e0206770 (2019).

10. K. Paap, H. Myuz, O. Sawi, Are bilingual advantages dependent upon specific tasks or specific bilingual experiences? Journal of Cognitive Psychology 26 (2014).

11. E. S. Nichols, C. J. Wild, B. Stojanoski, M. E. Battista, A. M. Owen, Bilingualism Affords No General Cognitive Advantages: A Population Study of Executive Function in 11,000 People. Psychol Sci 31, 548–567 (2020).

12. M. van den Noort, E. Struys, P. Bosch, Individual Variation and the Bilingual Advantage—Factors that Modulate the Effect of Bilingualism on Cognitive Control and Cognitive Reserve. Behav Sci (Basel) 9, 120 (2019).

13. A. S. Dick, et al., No evidence for a bilingual executive function advantage in the nationally representative ABCD study. Nat Hum Behav 3, 692–701 (2019).

14. A. de Bruin, A. S. Dick, M. Carreiras, Clear Theories Are Needed to Interpret Differences: Perspectives on the Bilingual Advantage Debate. Neurobiology of Language 2, 433–451 (2021).

15. J. A. O’Sullivan, et al., Attentional Selection in a Cocktail Party Environment Can Be Decoded from Single-Trial EEG. Cereb Cortex 25, 1697–1706 (2015).

16. S. A. Fuglsang, T. Dau, J. Hjortkjær, Noise-robust cortical tracking of attended speech in real-world acoustic scenes. NeuroImage 156, 435–444 (2017).

17. V. Commuri, J. P. Kulasingham, J. Z. Simon, Cortical responses time-locked to continuous speech in the high-gamma band depend on selective attention. Front. Neurosci. 17 (2023).

18. E. M. Zion Golumbic, et al., Mechanisms underlying selective neuronal tracking of attendedspeech at a “cocktail party.” Neuron 77, 980–991 (2013).

19. H. Ershaid, et al., Contributions of listening effort and intelligibility to cortical tracking of speech in adverse listening conditions. Cortex 172, 54–71 (2024).

20. B. D. Zinszer, Q. Yuan, Z. Zhang, B. Chandrasekaran, T. Guo, Continuous speech tracking in bilinguals reflects adaptation to both language and noise. Brain and Language 230, 105128 (2022).

21. M. Lizarazu, M. Carreiras, M. Bourguignon, A. Zarraga, N. Molinaro, Language proficiency entails tuning cortical activity to second language speech. Cerebral Cortex 31, 3820–3831 (2021).

22. G. M. Di Liberto, et al., Neural representation of linguistic feature hierarchy reflects second-language proficiency. NeuroImage 227, 117586 (2021).

23. A. S. Ihara, et al., Prediction of second language proficiency based on electroencephalographic signals measured while listening to natural speech. Front. Hum. Neurosci. 15, 665809 (2021).

24. R. Reetzke, G. N. Gnanateja, B. Chandrasekaran, Neural tracking of the speech envelope is differentially modulated by attention and language experience. Brain and Language 213, 104891 (2021).

25. J. Pérez-Navarro, et al., Early language experience modulates the tradeoff between acoustic-temporal and lexico-semantic cortical tracking of speech. iScience 27, 110247 (2024).

26. F. Tezcan, H. Weissbart, A. E. Martin, A tradeoff between acoustic and linguistic feature encoding in spoken language comprehension. eLife 12, e82386 (2023).

27. L. Van Den Eynde, M. Gillis, E. Rombouts, I. Zink, M. Vandermosten, Measuring language proficiency in bilingual children using EEG-based neural tracking of continuous speech. Brain and Language 280, 105801 (2026).

28. M. J. Crosse, et al., Linear Modeling of Neurophysiological Responses to Speech and Other Continuous Stimuli: Methodological Considerations for Applied Research. Front. Neurosci. 15 (2021).

29. M. J. Crosse, G. M. Di Liberto, A. Bednar, E. C. Lalor, The Multivariate Temporal Response Function (mTRF) Toolbox: A MATLAB Toolbox for Relating Neural Signals to Continuous Stimuli. Front. Hum. Neurosci. 10 (2016).

30. E. C. Lalor, A. J. Power, R. B. Reilly, J. J. Foxe, Resolving precise temporal processing properties of the auditory system using continuous stimuli. Journal of neurophysiology 102, 349–359 (2009).

31. V. Marian, H. K. Blumenfeld, M. Kaushanskaya, The Language Experience and Proficiency Questionnaire (LEAP-Q): assessing language profiles in bilinguals and multilinguals. J Speech Lang Hear Res 50, 940–967 (2007).

32. E. Ahissar, et al., Speech comprehension is correlated with temporal response patterns recorded from auditory cortex. Proceedings of the National Academy of Sciences 98, 13367–13372 (2001).

33. E. C. Lalor, J. J. Foxe, Neural responses to uninterrupted natural speech can be extracted with precise temporal resolution. European journal of neuroscience 31, 189–193 (2010).

34. N. Ding, J. Z. Simon, Emergence of neural encoding of auditory objects while listening to competing speakers. Proc Natl Acad Sci U S A 109, 11854–11859 (2012).

35. G. M. Di Liberto, J. A. O’Sullivan, E. C. Lalor, Low-Frequency Cortical Entrainment to Speech Reflects Phoneme-Level Processing. Current Biology 25, 2457–2465 (2015).

36. I. D. Karunathilake, J. P. Kulasingham, J. Z. Simon, Neural tracking measures of speech intelligibility: Manipulating intelligibility while keeping acoustics unchanged. Proceedings of the National Academy of Sciences 120, e2309166120 (2023).

37. A.-L. Giraud, D. Poeppel, Cortical oscillations and speech processing: emerging computational principles and operations. Nat Neurosci 15, 511–517 (2012).

38. N. Ding, J. Z. Simon, Cortical entrainment to continuous speech: functional roles and interpretations. Frontiers in Human Neuroscience Volume 8-2014 (2014).

39. N. Mesgarani, E. F. Chang, Selective cortical representation of attended speaker in multi-talker speech perception. Nature 485, 233–236 (2012).

40. J. E. Peelle, J. Gross, M. H. Davis, Phase-Locked Responses to Speech in Human Auditory Cortex are Enhanced During Comprehension. Cerebral Cortex 23, 1378–1387 (2013).

41. J. Vanthornhout, L. Decruy, J. Wouters, J. Z. Simon, T. Francart, Speech intelligibility predicted from neural entrainment of the speech envelope. Journal of the Association for Research in Otolaryngology 19, 181–191 (2018).

42. J. R. Kerlin, A. J. Shahin, L. M. Miller, Attentional gain control of ongoing cortical speech representations in a “cocktail party.” Journal of Neuroscience 30, 620–628 (2010).

43. D. Lesenfants, T. Francart, The interplay of top-down focal attention and the cortical tracking of speech. Sci Rep 10, 6922 (2020).

44. M. Kalashnikova, V. Peter, G. M. Di Liberto, E. C. Lalor, D. Burnham, Infant-directed speech facilitates seven-month-old infants’ cortical tracking of speech. Sci Rep 8, 13745 (2018).

45. L. D. A. Quan, L. T. Trang, I. Choi, J. Woo, Differential electroencephalography responses in speech perception between native and non-native speakers. Front. Hum. Neurosci. 19 (2025).

46. J. Zou, et al., Auditory and language contributions to neural encoding of speech features in noisy environments. NeuroImage 192, 66–75 (2019).

47. C. Daube, R. A. Ince, J. Gross, Simple acoustic features can explain phoneme-based predictions of cortical responses to speech. Current Biology 29, 1924–1937. e9 (2019).

48. G. M. Di Liberto, J. A. O’Sullivan, E. C. Lalor, Low-Frequency Cortical Entrainment to Speech Reflects Phoneme-Level Processing. Current Biology 25, 2457–2465 (2015).

49. E. S. Teoh, M. S. Cappelloni, E. C. Lalor, Prosodic pitch processing is represented in delta-band EEG and is dissociable from the cortical tracking of other acoustic and phonetic features. European Journal of Neuroscience 50, 3831–3842 (2019).

50. M. P. Broderick, A. J. Anderson, G. M. D. Liberto, M. J. Crosse, E. C. Lalor, Electrophysiological Correlates of Semantic Dissimilarity Reflect the Comprehension of Natural, Narrative Speech. Current Biology 28, 803–809.e3 (2018).

51. M. Heilbron, K. Armeni, J.-M. Schoffelen, P. Hagoort, F. P. De Lange, A hierarchy of linguistic predictions during natural language comprehension. Proceedings of the National Academy of Sciences 119, e2201968119 (2022).

52. P. Boersma, D. Weenink, Praat: Doing phonetcs by computer. (2022). Deposited 2022.

53. The MathWorks, Inc., MATLAB R2022a (Version 2022a). (2022). Deposited 2022.

54. S. H. Jessica Tan, M. Kalashnikova, G. M. Di Liberto, M. J. Crosse, D. Burnham, Seeing a talking face matters: The relationship between cortical tracking of continuous auditory-visual speech and gaze behaviour in infants, children and adults. NeuroImage 256, 119217 (2022).

55. Posit Software, PBC, RStudio: Integrated Development Environment for R. (2025). Deposited 2025.

